# Epigenomic signatures as circulating and predictive biomarkers in sarcomatoid renal cell carcinoma

**DOI:** 10.1101/2023.11.27.568827

**Authors:** Talal El Zarif, Karl Semaan, Marc Eid, Ji-Heui Sep, Simon Garinet Semaan, Matthew P. Davidsohn, Brad Fortunato, John Canniff, Amin H. Nassar, Sarah Abou Alaiwi, Ziad Bakouny, Gitanjali Lakshmi, Hunter Savigano, Kevin Lyons, Sayed Matar, Atef Ali, Eddy Saad, Renee Maria Saliby, Paulo Cordeiro, Ziwei Zhang, Nourhan El Ahmar, Yasmine Nabil Laimon, Chris Labaki, Valisha Shah, Dory Freeman, Jillian O’Toole, Gwo-Shu Mary Lee, Justin Hwang, Mark Pomerantz, Sabina Signoretti, Wanling Xie, Jacob E. Berchuck, Srinivas R. Viswanathan, Eliezer M. Van Allen, David A. Braun, Toni K. Choueiri, Matthew L. Freedman, Sylvan C. Baca

**Affiliations:** Department of Internal Medicine, Yale School of Medicine, New Haven, CT, USA; Department of Medical Oncology, Dana-Farber Cancer Institute; Boston, MA, USA; Section of Medical Oncology, Department of Internal Medicine, Yale School of Medicine, New Haven, CT, USA; Section of Cardiology, Department of Internal Medicine, Yale School of Medicine, New Haven, CT, USA; Department of Medical Oncology, Memorial Sloan Kettering Cancer Center, New York, NY, USA; Department of Pathology, Brigham and Women’s Hospital, Boston, Massachusetts, USA; Department of Medicine, University of Minnesota Masonic Cancer Center, Minneapolis, MN, USA

## Abstract

Renal cell carcinoma with sarcomatoid differentiation (sRCC) is associated with poor survival and heightened response to immune checkpoint inhibitors (ICIs). Two major barriers to improving outcomes for sRCC are (1) a limited understanding of its gene regulatory programs and (2) difficulty identifying sarcomatoid differentiation on tumor biopsies due to spatial heterogeneity. To address these challenges, we characterized the epigenomic landscape of sRCC by profiling 107 epigenomic libraries in tissue and plasma samples from 50 patients with RCC and healthy volunteers. We identified highly recurrent epigenomic reprogramming, as assessed by histone modifications and DNA methylation, that distinguishes sRCC from non-sarcomatoid RCC. Computational analysis of RCC epigenomic profiles and CRISPRa experiments implicated the transcription factor FOSL1 in activating sRCC-associated gene regulatory programs. Analysis of two randomized clinical trials identified FOSL1 expression as a predictive biomarker of response to ICIs in RCC. Finally, we demonstrate that epigenomic signatures of sRCC are detectable in patient plasma, establishing an approach for blood-based diagnosis of this clinically important phenotype. These findings provide a framework for the discovery and non-invasive detection of epigenomic correlates of tumor histology via liquid biopsy.

## Introduction

Sarcomatoid differentiation is a histopathologic feature observed in 5-20% of renal cell carcinoma (RCC) tumors across all histologies^1^, including clear cell RCC (ccRCC), the most common subtype of kidney cancer^2,3^. RCC with sarcomatoid features (sRCC) exhibits an aggressive phenotype characterized by epithelial-to-mesenchymal transition (EMT) and responds poorly to vascular endothelial growth factor receptor tyrosine kinase inhibitors (VEGFR TKIs) compared to non-sarcomatoid or “epithelioid” RCC^4–6^. Diagnosing sRCC is clinically important because it responds particularly well to immune checkpoint inhibitors (ICIs)^6–10^, potentially due to increased PD-L1 expression on tumor cells and prominent immune cell infiltration^6,10–13^.

The molecular underpinnings of sarcomatoid differentiation are poorly understood, partly due to a paucity of cell line models that faithfully recapitulate sRCC biology^10^. Recent studies have therefore focused on the analysis of clinical sRCC tissues to identify genomic and transcriptomic features that may drive the aggressive behavior of sRCC. Compared to epithelioid ccRCC, sRCC is enriched for several mutations and transcriptional features, such as deletions of *CDKN2A*/*CDKN2B*^10^ and increased expression of cell cycle genes (i.e., E2F7)^6^. Yet, these cancers are not distinguishable based on mutational profiles. ccRCC and sRCC components from the same patient generally share mutations, suggesting a common clonal origin^10,14–17^. In contrast to transcriptional studies, data on the epigenomic programs that drive EMT in sRCC are lacking^1^.

Sarcomatoid differentiation is spatially heterogeneous, and sRCC and epithelioid RCC components often co-exist within a tumor. Therefore, the detection of sarcomatoid differentiation using tumor biopsy is challenging. In two case series, biopsies detected sarcomatoid features in only 7.5% and 9.1% of patients who were later found to have clear evidence of sRCC in their nephrectomy specimens^18,19^. This limitation is compounded by the fact that patients with metastatic RCC are increasingly diagnosed from single tissue biopsies as cytoreductive nephrectomies have fallen out of favor in metastatic RCC^20,21^. Therefore, tractable biomarkers that reflect the aggregate burden of cancer are needed for sRCC diagnoses. To this end, recent studies have shown promise in detecting clonally related histologic variants using epigenomic signatures from plasma^22–24^.

In this study, we characterized the epigenomes of sarcomatoid and epithelioid ccRCC tumors from pathologically reviewed clinical tissue specimens. We hypothesized that sarcomatoid differentiation is driven by the activation of *cis*-regulatory elements (CREs) bound by a set of master transcription factors (TFs). We identified marked and consistent differences in the epigenomic profiles of sRCC compared to epithelioid ccRCC. Based on differential regulatory elements, we nominated candidate TFs that may drive epigenomic changes that characterize sRCC. We found that TFs implicated by epigenomic profiling are upregulated in sRCC across three clinical cohorts and that their expression levels are associated with clinical outcomes. We experimentally confirmed that the expression of a candidate TF in an epithelioid ccRCC cell line model activates sarcomatoid-like gene expression patterns. Finally, we applied tissue-derived epigenomic signatures to detect sRCC in plasma using epigenomic liquid biopsy techniques (Figure 1). Overall, our findings uncover epigenetic changes in gene regulation that may drive sarcomatoid differentiation in RCC and suggest an approach for blood-based detection of sarcomatoid differentiation to guide therapy selection.

**Figure 1.**
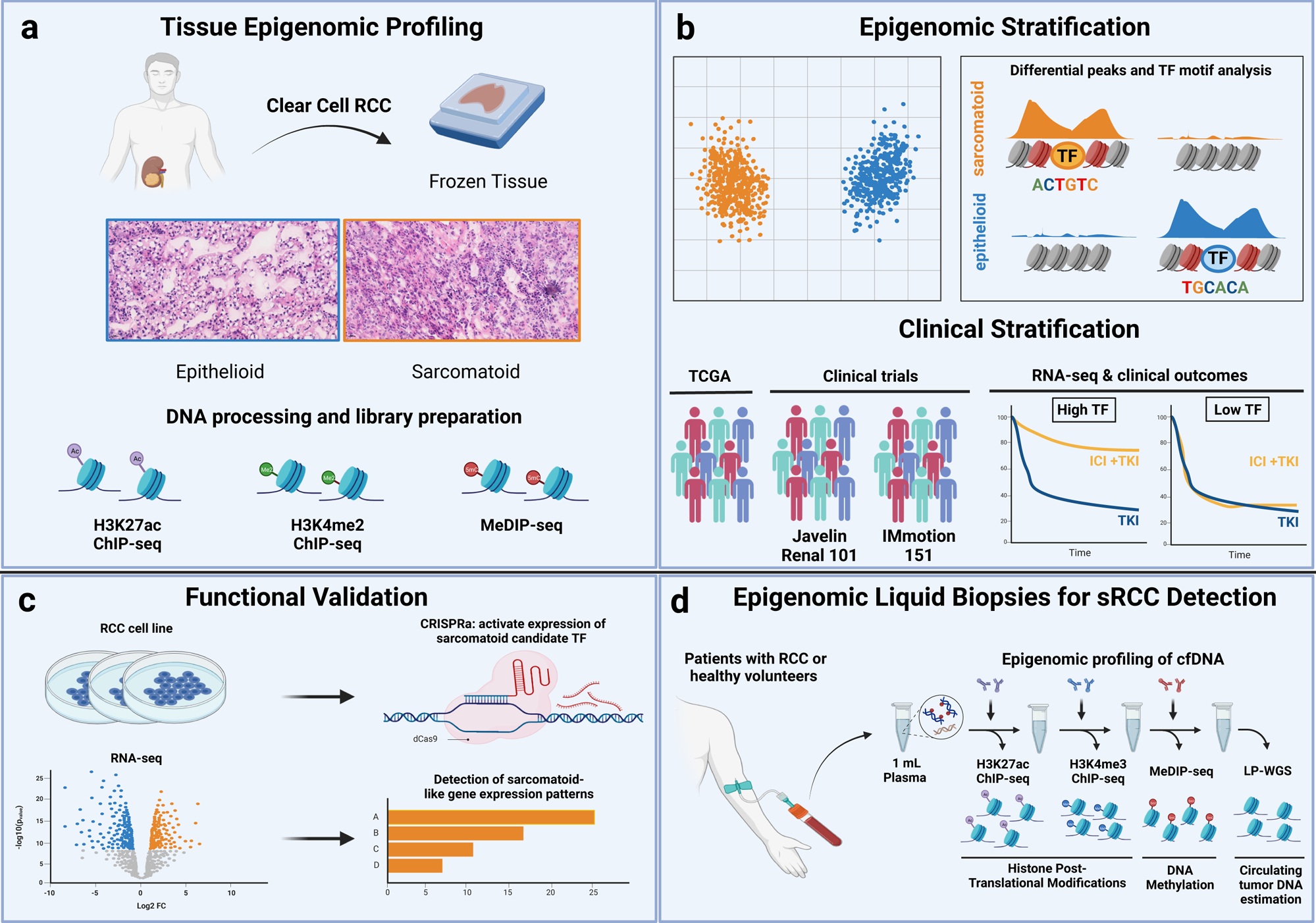
Tissue- and plasma-based epigenomic characterization of sRCC. **a** Epigenomic profiling of frozen tissue samples from patients with epithelioid or sarcomatoid ccRCC. **b** Stratification of tissue samples by subtype-specific epigenomic features and their correlation with clinical outcomes in three cohorts. **c** Detection of sarcomatoid-like gene expression patterns following activation of expression of a candidate transcription factor in an RCC cell line model. **d** Plasma-based epigenomic profiling of patients with ccRCC and healthy volunteers. ccRCC: clear cell renal cell carcinoma, TF: transcription factor: ICI: immune checkpoint inhibitors, TKI: tyrosine kinase inhibitors, LP-WGS: low pass whole genome sequencing, cfDNA: cell-free DNA.

## Results

We generated 107 tissue- and plasma-based epigenomic libraries from 50 individuals across different epigenomic assays. Pathologically reviewed tissue samples were derived from 31 individual patients with ccRCC (10 sarcomatoid and 21 epithelioid). Plasma was collected from 19 individuals: 5 with sRCC, 5 with epithelioid ccRCC, and 9 healthy volunteers.

### Sarcomatoid and epithelioid ccRCC exhibit distinct epigenomic profiles

Using tissue samples, we examined the epigenomic landscape of sRCC (Supplementary Data 1, Fig. 2A). We performed chromatin immunoprecipitation sequencing (ChIP-seq) for post-translational histone modifications (H3K27ac and H3K4me2) and methylated CpG dinucleotide sequencing (MeDIP-seq). H3K27ac is associated with active promoters and enhancers^25^, while H3K4me2 ChIP-seq mostly reflects active and poised promoters, in addition to enhancers^26,27^. Peaks for each epigenetic mark overlapped expected genomic annotations. For example, 61% of H3K27ac peaks overlapped with non-promoter regions of annotated genes, consistent with the capture of promoter-distal active enhancers (Fig. 2B). H3K27ac and H3K4me2 ChIP-seq signals overlapped different genomic annotations, with H3K4me2 capturing more promoters and H3K27ac capturing more promoter-distal CREs (Fig. 2B), which is in line with prior reports^28–30^. Principal component analyses (PCA) (Fig. 2C, D, E) and unsupervised hierarchical clustering (Supplementary Fig. S1A, B, C) of H3K27ac and H3K4me2 peaks and methylated CpG islands cleanly separated epithelioid and sRCC samples. This highlighted consistent differences in the epigenomic features between groups, with 25,919 H3K27ac, 44,133 H3K4me2, and 51,464 MeDIP peaks upregulated in sRCC or epithelioid ccRCC (q < 0.05, Fig. 2F).

**Figure 2.**
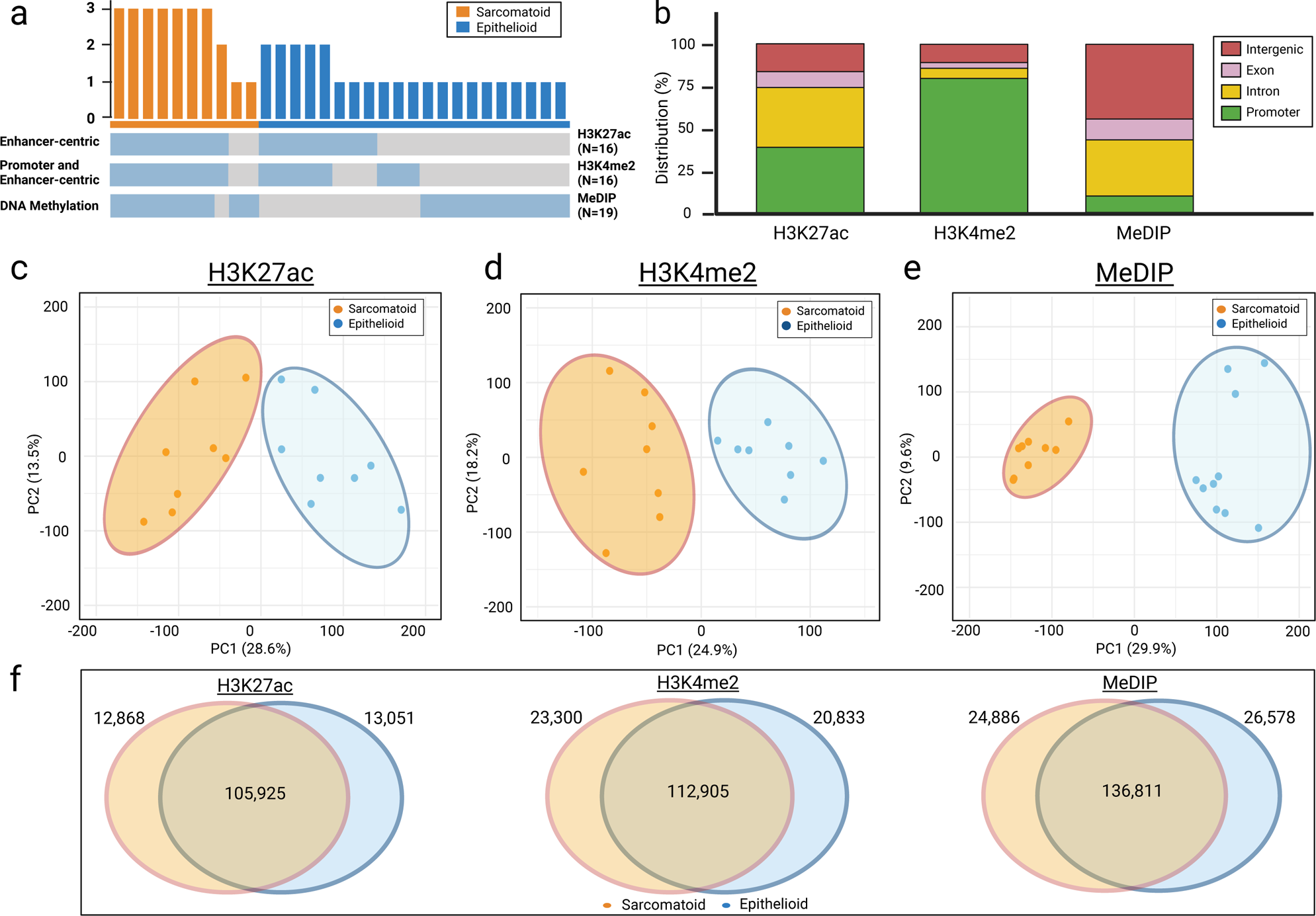
H3K27ac, H3K4me2, and methylated CpG dinucleotide regions between sarcomatoid and epithelioid ccRCC reflect distinct epigenomic programs. **a** Epigenomic datasets generated from tissue samples. **b** Distribution of H3K27ac, H3K4me2, and MeDIP peaks by genomic regions **c-e** Principal component analysis (PCA) plots of the H3K27ac, H3K4me2, and MeDIP peaks in epithelioid and sarcomatoid ccRCC. **f** Venn-Diagrams of the subtype-enriched H3K27ac, H3K4me2, and MeDIP peaks.

H3K27ac enables the identification of active programs of gene regulation by capturing active promoters and enhancers. To evaluate the biological significance of sRCC- and epithelioid-enriched H3K27ac CREs (Fig. 3A), we evaluated the enrichment of gene ontology (GO) terms of genes near differential CREs using GREAT^31^. We noted significant enrichment at the top 1,621 sRCC CREs (q < 0.01, LFC > 1) for gene pathways involved in immune cell activation and stimulation of an immune response (Fig. 3B). These results are consistent with previous reports demonstrating that sRCC tumors show increased inflammatory gene expression signatures and CD8+ T cell infiltration, and have higher expression of PD-1/L1^10,13^, which may partly explain their responsiveness to ICIs^6–8,10^. Additionally, we observed processes implicated in shifts in cellular morphology and extracellular matrix organization, which are expected in the setting of EMT, a defining feature of sarcomatoid differentiation^32^. A similar analysis of the top 3,386 epithelioid-specific H3K27ac CREs (q < 0.01, LFC > 1) showed enrichment for genes involved in the response to hypoxia (Fig. 3C), likely reflecting the activity of hypoxia-inducible factors (i.e., HIF2α) that are implicated in angiogenesis and RCC carcinogenesis^33^. These findings are consistent with observations from the IMmotion151 (IM151) trial^34^, where sRCC tumors exhibited more immunogenic and less angiogenic gene expression profiles than their epithelioid counterparts^6^.

**Figure 3.**
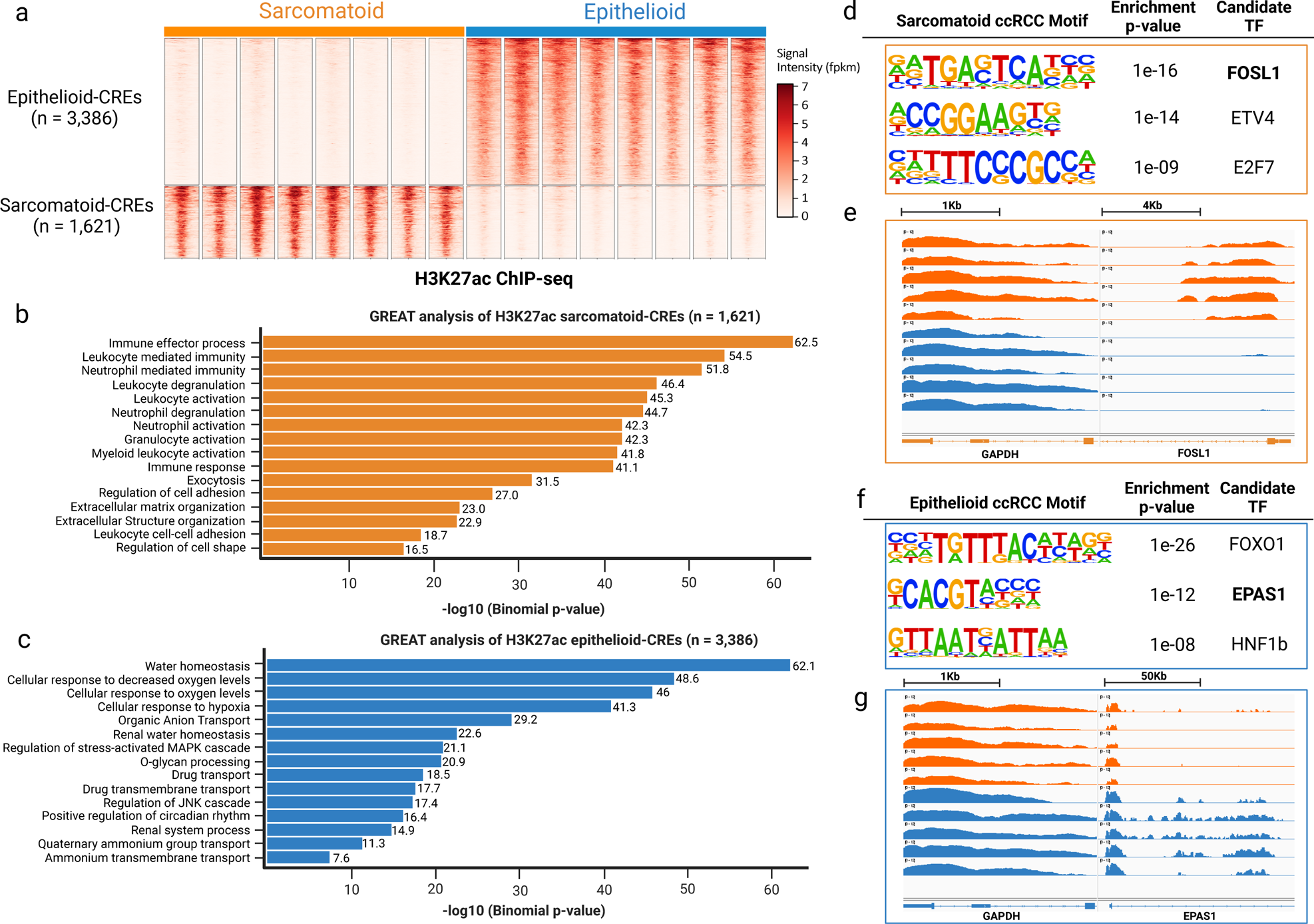
Sarcomatoid-enriched CREs are associated with immune system activation and drivers of EMT. **a** Heatmaps of normalized H3K27ac tag densities at differential H3K27ac *cis-*regulatory elements (CREs) between epithelioid and sarcomatoid ccRCC samples located ±2 kb from peak center. **b** GREAT analysis of sarcomatoid-enriched H3K27ac peaks (*n* = 1,621). **c** GREAT analysis of epithelioid-enriched H3K27ac peaks (*n* = 3,386). **d** Three top non-redundant significantly enriched nucleotide motifs present in sarcomatoid-specific sites by de novo motif analysis. **e** H3K27ac profiles near *FOSL1* in five representative samples of each ccRCC subtype, normalized to signal at GAPDH. **f** Three significantly enriched nucleotide motifs present in epithelioid-specific sites by de novo motif analysis. **g** H3K27ac profiles near *EPAS1* normalized to *GAPDH* in five representative samples of each ccRCC subtype (epithelioid and sarcomatoid).

Next, we analyzed differential H3K27ac CREs (q < 0.01, LFC > 1) for the enrichment of nucleotide motifs to identify TFs that may drive CRE activation. The top three motifs that were highly enriched at sarcomatoid-specific CREs were FOSL1, ETV4, and E2F7 (Fig. 3D). The *FOSL1* gene demonstrated a higher H3K27ac signal in sarcomatoid vs. epithelioid ccRCC samples (Fig. 3E). FOSL1 is a TF and a subunit of the AP-1 complex implicated in invasiveness and EMT in prostate^35^, breast^36–38^, and kidney cancers^39^. Conversely, motif enrichment analysis of putative enhancers downregulated in sRCC identified FOXO1, EPAS1, and HNF1β as the top candidate TFs (Fig. 3F)*. EPAS1*, which encodes HIF2α, is densely marked with H3K27ac in epithelioid vs. sRCC and is a central TF in ccRCC oncogenesis (Fig. 3G). Overall, differential CREs pointed to biologically plausible TFs that may activate EMT programs in sRCC and implicated decreased activity of developmental (HNF1β) and hypoxia-related (HIF2α) TFs in sarcomatoid differentiation.

### Candidate sarcomatoid TFs predict clinical outcomes independent of histologic subtyping

Having nominated TFs that may drive EMT-associated gene regulatory programs of sRCC, we tested whether their expression levels in RCC are associated with clinical outcomes. We analyzed transcriptomic data from TCGA^2,3^ and two large phase III randomized clinical trials: Javelin Renal 101 (JR101)^40^ and IM151 cohorts. In JR101, patients were randomized to the avelumab plus axitinib combination or sunitinib arms, while in IM151, they were randomized to atezolizumab plus bevacizumab combination or sunitinib arms. We identified genes significantly upregulated or downregulated in sRCC tumors from the three cohorts (FDR q < 0.05) and selected the upregulated TFs. Strikingly, the top three TFs we nominated (FOSL1, E2F7, and ETV4) were upregulated in sRCC and were associated with improved progression-free survival (PFS) in patients with RCC who were treated with ICI plus TKI combinations vs. TKIs alone (Supplementary Data 2).

Next, we aimed to identify TFs that may serve as predictive biomarkers for ICIs in RCC. We focused on FOSL1 since it was the top-ranked TF in the motif analysis, had an evident increase in the H3K27ac signal at the *FOSL1* gene locus, and had the strongest association with shorter PFS in the clinical trials. *FOSL1* expression was higher in sRCC vs. epithelioid ccRCC in the TCGA KIRC cohort (q = 4.91×10^−11^, LFC = 1.73), JR101 (q = 8.51×10^−12^, LFC = 1.23), and IM151 (q = 2.11×10^−11^, LFC = 0.64) (Fig. 4A). In a multivariable analysis accounting for the IMDC risk groups, prognostic factors in RCC^4^, patients with high (> median) *FOSL1*-expressing tumors had improved PFS when treated with ICI plus TKI combinations vs. sunitinib in both JR101 (adjusted HR: 0.53, 95%CI: 0.41 – 0.70, p <0.001, Fig. 4B) and IM151 (adjusted HR: 0.71, 95%CI: 0.56 – 0.90, p = 0.004; Supplementary Fig. S2). Patients with low (< median) *FOSL1-* expressing tumors had similar PFS regardless of ICI use in both JR101 (adjusted HR: 0.85; 95%CI: 0.63 - 1.13; p = 0.25, Fig. 4B) and IM151 (adjusted HR: 1.02; 95%CI: 0.80 - 1.32, p = 0.86, Supp Fig. S2). We included an interaction term between treatment arms (ICI + TKI vs. TKI) and *FOSL1* expression (high vs. low) in the Cox regression analysis for PFS, with treatment arm and baseline *FOSL1* expression as two other independent variables. Compared to sunitinib, ICI + TKI combinations improved PFS for patients with high *FOSL1*-expressing tumors but offered no benefit for patients with low *FOSL1* in JR101 (interaction p-value = 0.03) and IM151 (interaction p-value = 0.04) (Figure 4B, Supplementary Data 2). These results indicate that *FOSL1* expression may be a predictive biomarker for prolonged PFS in patients receiving ICIs + VEGFR TKIs in RCC.

**Figure 4.**
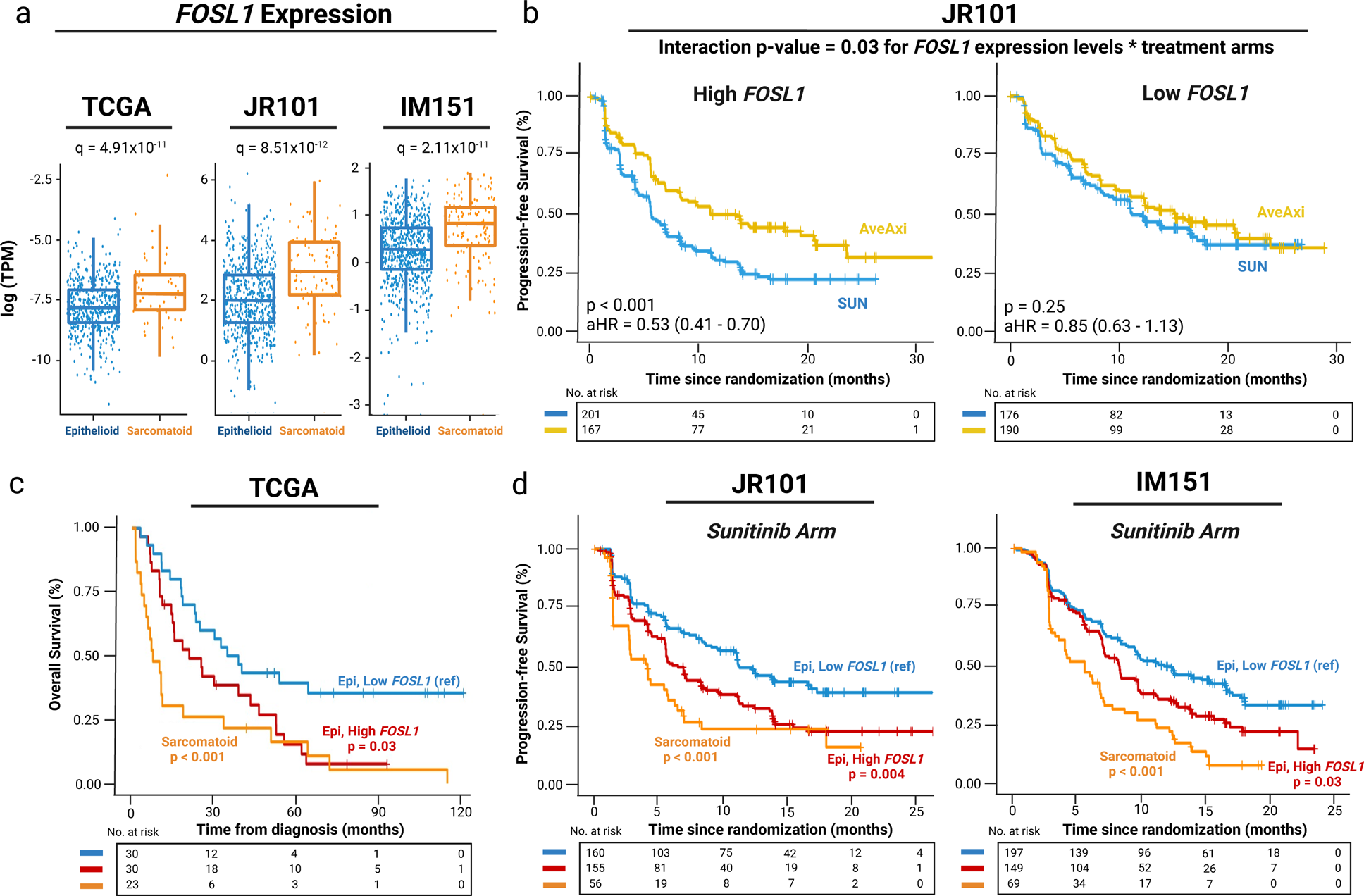
FOSL1 is upregulated in sRCC and is associated with clinical outcomes. **a** Boxplots of *FOSL1* expression levels in three clinical cohorts (TCGA, JR101, and IM151). **b** Progression-free survival in patients in JR101 by *FOSL1* levels. **c** Overall survival in patients with sRCC and epithelioid ccRCC divided by expression of *FOSL1* (High vs. Low) in the TCGA cohort. **d** Progression-free survival in patients with sRCC and epithelioid ccRCC divided by expression of *FOSL1* (High vs. Low) in the sunitinib arms of JR101 and IM151. TPM: Transcripts per million, TCGA: The Cancer Genome Atlas, JR101: Javelin Renal 101, IM151: Immotion151, Epi: epithelioid ccRCC, AveAxi: avelumab plus axitinib, AtezoBev: atezolizumab plus bevacizumab, SUN: sunitinib.

Notably, while *FOSL1* was upregulated in sRCC, many epithelioid ccRCC tumors also exhibited elevated *FOSL1* expression levels (Fig. 4A). In this context, we hypothesized that these tumors may behave similarly to sRCC, with worse outcomes with sunitinib compared to improved outcomes with ICI plus TKI combinations. Accordingly, we stratified patients into three groups: epithelioid ccRCC with low *FOSL1* or high *FOSL1*, and sRCC. Strikingly, the OS (time from diagnosis to death or loss of follow-up) of patients with epithelioid ccRCC and high *FOSL1* was worse compared to those with low *FOSL1* (median OS = 21.2 vs. 37.4 months; HR: 1.9; 95% CI: 1.1 – 3.5; p = 0.03) (Fig. 4C) and similar to those with sRCC tumors (p = 0.1) in the metastatic ccRCC TCGA cohort. Furthermore, patients with epithelioid ccRCC and high *FOSL1* had shorter PFS compared to those with low *FOSL1* in both JR101 (median PFS = 6.5 vs. 11.3 months; adjusted HR: 1.54; 95% CI: 1.14 – 2.06; p = 0.004) and IM151 (median PFS = 8.3 vs. 11.8 months; adjusted HR: 1.35; 95% CI: 1.03 – 1.76; p = 0.03, Fig. 4D), and similar PFS compared to sRCC tumors in the sunitinib arm of both trials (Supplementary Data 3). On the other hand, there were no significant differences in PFS between epithelioid ccRCC subgroups in the ICI arms, suggesting that all subgroups derived similar benefit (Supplementary Fig. S2). Taken together, the above findings indicate that elevated *FOSL1* expression is associated with worse outcomes when treated with TKI-only regimens, similar to sRCC, but outcomes may be improved with the addition of ICIs. These findings implicate *FOSL1* expression as a biomarker that recapitulates clinical outcomes in patients with ccRCC, independent of histopathologic subtyping.

### FOSL1 induces a sarcomatoid-like gene expression profile in epithelioid ccRCC

To test whether *FOSL1* drives processes observed in sRCC tumors (i.e., cell cycle progression^10,41^), we endogenously activated *FOSL1* expression in the Caki-1 ccRCC cell line using CRISPRa.

We evaluated the transcriptomic and epigenomic differences induced by activating *FOSL1* expression. We compared differentially expressed genes (q < 0.05) between triplicates of *FOSL1*-expressing vs. control Caki-1 cells (Methods, Fig. 5A). Overall, 357 genes were upregulated and 540 downregulated in *FOSL1* CRISPRa vs. control Caki-1 cells. As expected, *FOSL1* was upregulated in *FOSL1* CRISPRa cells (LFC = 0.59, q = 1.62×10^−10^), but strikingly, *EPAS1* was downregulated (LFC = −0.40, q = 9.95×10^−6^). The downregulation of *EPAS1* suggests that FOSL1 mediate the suppressed hypoxia and angiogenesis signaling observed in sRCC (Fig. 5B). Gene set enrichment analysis (GSEA) of *FOSL1* CRISPRa vs. negative control Caki-1 cells revealed upregulation of seven gene sets (Fig. 5C) with enrichment of programs related to cell cycle (E2F targets, G2M checkpoints) or proliferation (MYC targets), similar to prior observations^6,10^. Furthermore, the H3K27ac signal at the promoters of upregulated genes in CRISPRa *FOSL1* were increased compared to control Caki-1 cells, whereas they were decreased at the promoters of downregulated genes in CRISPRa *FOSL1* (Supplementary Fig. S3A). Taken together, the observed changes in gene expression correlated with biologically relevant changes in epigenomic H3K27ac signals and followed expected trends for both upregulated and downregulated genes in each condition.

**Figure 5.**
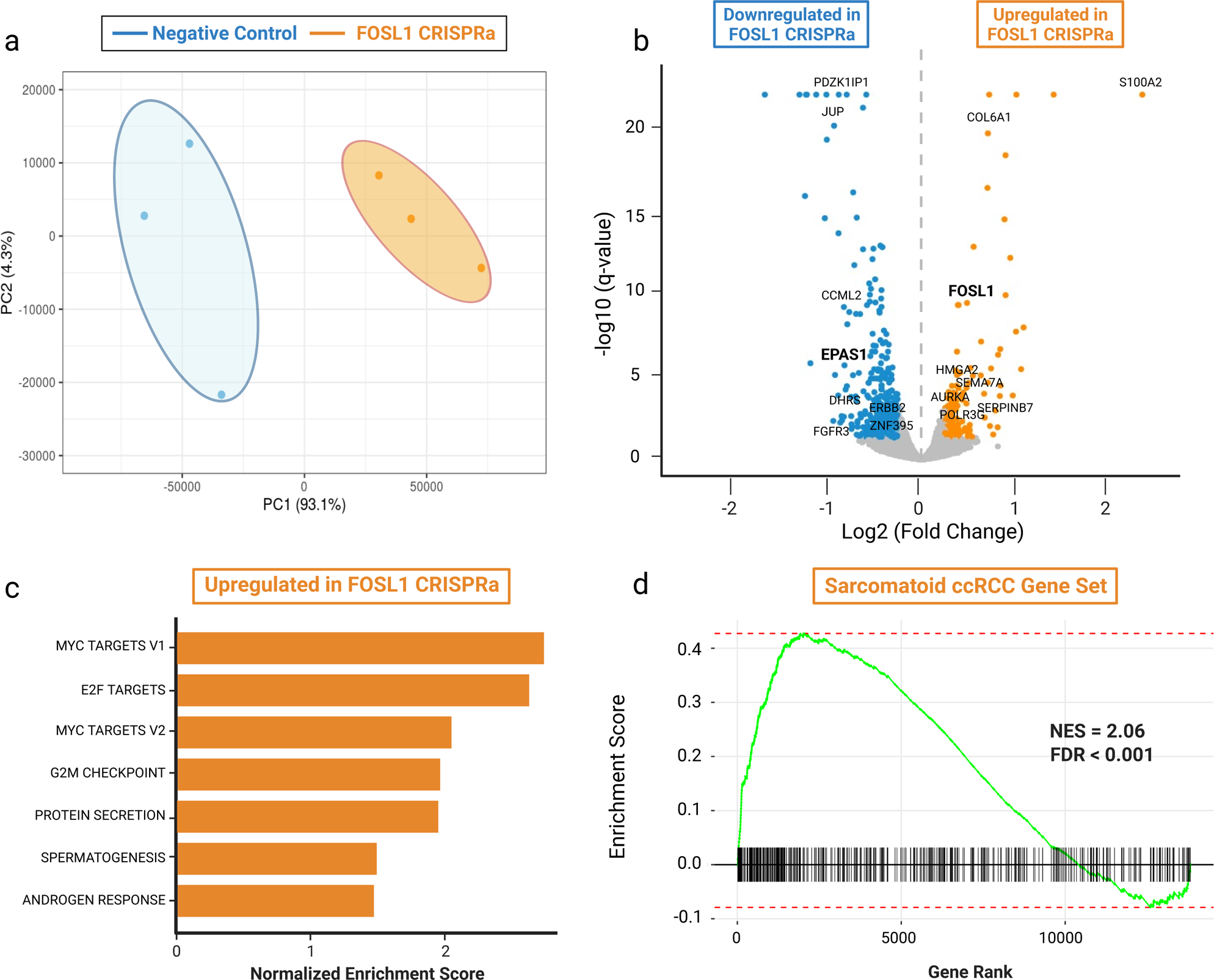
Expression of *FOSL1* in an epithelioid ccRCC cell line activates sRCC transcriptional programs. **a** Principal Component Analysis (PCA) plot of three replicates of RNA-sequencing data from two conditions (*FOSL1* CRISPRa and negative control Caki-1 cells). **b** Volcano plot showing upregulated and downregulated genes in *FOSL1* CRISPRa vs. negative control Caki-1 cells. **c** Normalized enrichment scores of pathways upregulated in *FOSL1* CRISPRa Caki-1 cells. **d** Gene set enrichment analysis (GSEA) of genes upregulated in sarcomatoid ccRCC from JR101 in *FOSL1* CRISPRa Caki-1 cells. ccRCC: clear cell renal cell carcinoma, NES: normalized enrichment score, FDR: false-detection rate. JR101: Javelin Renal 101.

We next validated our findings in an external cohort. Using gene expression data from JR101 trial, we defined a set of the top 500 genes that were upregulated in sRCC and a set of the top 500 genes upregulated in epithelioid RCC tumors. GSEA revealed that the sarcomatoid gene set is enriched among upregulated genes in *FOSL1* CRISPRa vs. control Caki-1 cells with a normalized enrichment score (NES) = 2.06, q <0.001 (Fig. 5D). The epithelioid ccRCC gene set is similarly enriched among the downregulated genes (NES = −1.43, q = 0.01, Supplementary Fig. S3B). In summary, these data demonstrate that overexpression of a single TF (FOSL1) upregulates transcriptional programs observed in sRCC.

### Epigenomic signatures of sRCC are detectable from circulating plasma nucleosomes

Detecting sarcomatoid features in RCC tumors has therapeutic implications since they predict poor response to TKIs and heightened response to ICIs. sRCC detection is challenging due to spatial heterogeneity and potential sampling errors, likely leading to substantial underdiagnosis^13^. To overcome this challenge, we investigated whether we could detect epigenomic features of sRCC in patient plasma as a proxy for histopathologic sarcomatoid features. Using a recently described liquid biopsy approach for epigenomic profiling from 1 mL of plasma^24^, we profiled H3K4me3, H3K27ac and DNA Methylation in plasma samples from an independent group of patients with sarcomatoid ccRCC (n=5), epithelioid ccRCC (n=5), and healthy volunteers without a history of cancer (n=9) (Supplementary Data 4, Fig. 6A, B). We inferred the circulating tumor DNA (ctDNA) content in plasma samples by applying ichorCNA to LPWGS data^42^. The average (standard deviation) estimated ctDNA content was 0.024 (± 0.022) in plasma samples from patients with RCC (n=10) and 0 (± 0) in plasma samples of healthy volunteers (n=9) (Supplementary Data 4). Plasma H3K4me3 signal was elevated at the *PAX8* gene locus in patients with RCC compared to healthy volunteers, regardless of the presence or absence of sarcomatoid features (Fig. 6C), indicating that we are sampling epigenomic signals from RCC cells. PAX8 is a histopathologic marker for RCC^43^ that is uniquely expressed in RCC tumors and is involved in kidney embryogenesis^44^ and RCC. Furthermore, plasma H3K27ac signal was elevated in RCC patient plasma at RCC-specific regulatory elements previously defined in TCGA tumors based on chromatin accessibility^45^. These findings confirmed that our plasma-based epigenomic assay can detect RCC-specific signals.

**Figure 6.**
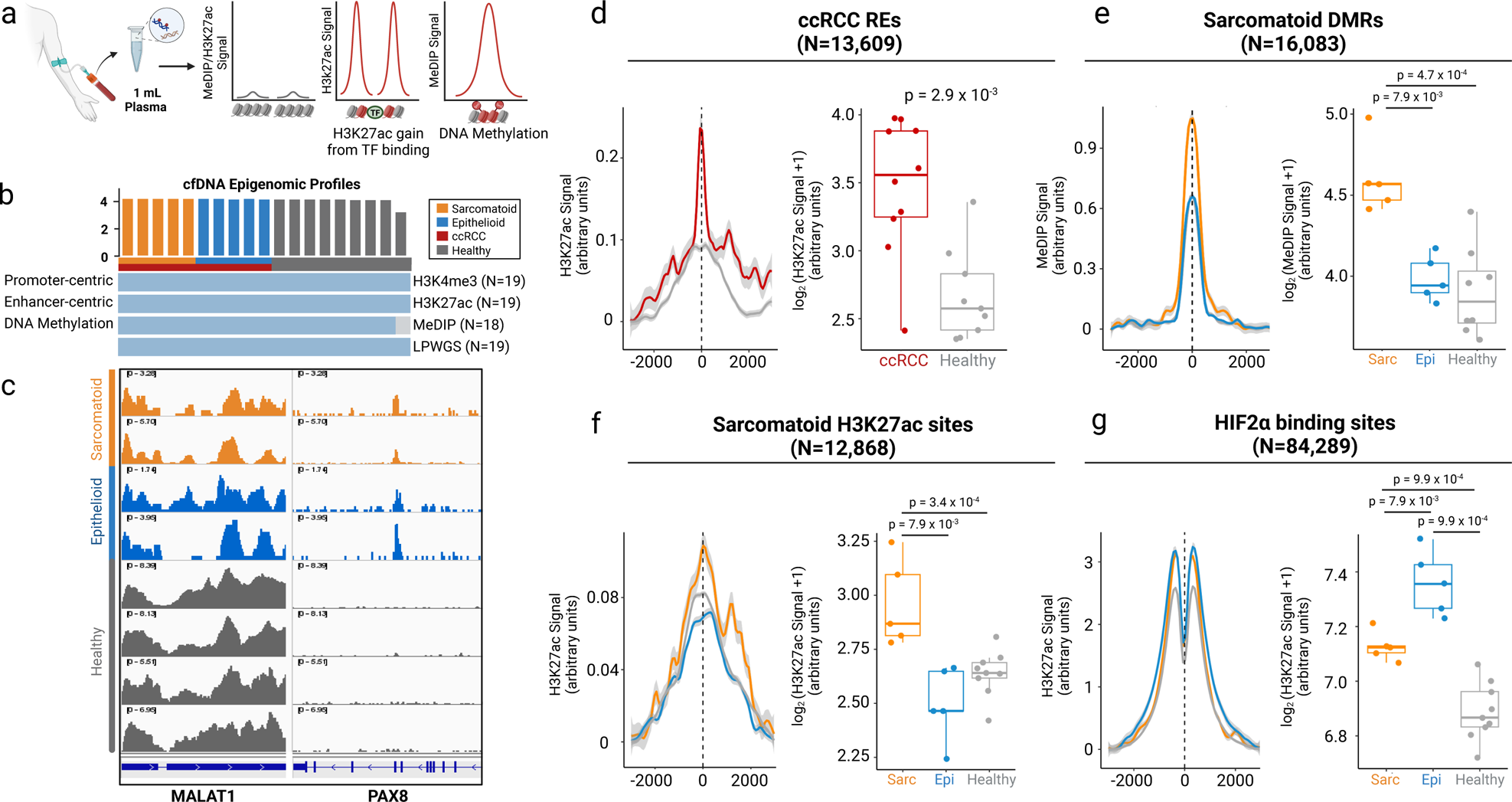
Tissue-informed epigenomic signatures enable the detection of sRCC in plasma. **a** Schematic demonstrating the measurement of cfChIP and cfMeDIP signals at TF binding sites differentially methylated regions, respectively. **b** Epigenomic datasets generated from plasma. **c** H3K4me3 cfChIP profiles at PAX8 in plasma of patients with sRCC (n=2), epithelioid ccRCC (n=2) and healthy volunteers (n=4). **d** Aggregated H3K27ac cfChIP-seq signals in kidney cancer and healthy plasma at REs identified by ATAC-seq in ccRCC. **e** Aggregated cfMeDIP-seq signals at tissue-informed upregulated sarcomatoid-specific DMRs and comparison between sRCC, epithelioid ccRCC, and healthy plasma, respectively. **f** Aggregated H3K27ac cfChIP-seq signals at tissue-informed upregulated sarcomatoid-specific H3K27ac sites and comparison between sRCC, epithelioid ccRCC, and healthy plasma, respectively. **g** Aggregated H3K27ac cfChIP-seq signal at HIF2α binding sites for ccRCC and comparison between sRCC, epithelioid ccRCC, and healthy plasma, respectively. **d-g** Dark lines show the mean signal across all samples in the indicated class. Gray shades represent the standard errors of the means. Boxplots indicate the area under the curve for the aggregate H3K27ac or MeDIP profiles for each sample. Wilcoxon test p-values are indicated for comparison across the groups: ccRCC (red), healthy (gray), epithelioid ccRCC (blue), and sRCC (orange). ccRCC: clear cell renal cell carcinoma, Sarc: sarcomatoid ccRCC (sRCC), Epi: epithelioid ccRCC, RE: regulatory elements, DMR: differentially methylated regions.

Next, we evaluated whether we could capture sarcomatoid-specific epigenomic features in plasma. We focused on the set of sarcomatoid-enriched DMRs in tissue samples to inform cfMeDIP-seq in plasma. We found that there were higher aggregate methylation values at these sites in sRCC plasma samples compared to plasma samples of patients with epithelioid ccRCC (p = 7.9 x 10^−3^) or healthy volunteers (p = 4.7 x 10^−4^, Fig. 6E). Similarly, the H3K27ac cfChIP-seq signal was increased at differential H3K27ac sites derived from tissue samples in plasma of patients with sRCC vs. those with epithelioid ccRCC tumors (p = 7.9 x 10^−3^) or healthy volunteers (p = 3.4 x 10^−4^, Fig. 6F). Importantly, we observed increased H3K27ac in plasma at binding sites for HIF2α in epithelioid tumors compared to sRCC (p = 7.9 x 10^−3^) and healthy plasma (p = 9.9 x 10^−4^, Fig. 6G). These findings align with our observations from the epigenomic and transcriptomic datasets, showing that sarcomatoid differentiation involves down-regulation of *EPAS1* and HIF2α activity. Overall, these results serve as proof of concept for detecting sRCC in plasma using minimally invasive liquid biopsy techniques.

## Discussion

In this study, we present the first epigenomic characterization of sRCC from patient tissues and plasma. We demonstrate that highly recurrent *cis*-regulatory programs are activated in sRCC compared to epithelioid ccRCC. These regulatory programs likely drive differential gene expression that has previously been reported between these groups^10,15^. Identification of sRCC-enriched CREs enabled us to nominate candidate TFs that drive sRCC gene regulatory programs. Using data from TCGA and two clinical trials, we show that these TFs are upregulated in sRCC and that their expression levels are associated with worse clinical outcomes. We identify *FOSL1* expression as a driver of sRCC and a predictive biomarker in RCC. Finally, we identify epigenomic fingerprints of sRCC in plasma based on circulating DNA methylation and histone modifications.

Our findings provide biological insights into the development of sRCC by nominating key TFs that may orchestrate transcriptional programs associated with sarcomatoid differentiation. FOSL1 emerged as a top candidate master TF from motif enrichment analysis of sarcomatoid CREs. FOSL1 is a subunit of AP-1 involved in tumor cell proliferation, survival, and invasiveness^46^ that may collaborate with ETV4, another candidate sRCC TF, to promote EMT and metastasis in RCC^39^. Importantly, activation of *FOSL1* expression in an epithelial ccRCC cell line decreased expression of *EPAS1* and upregulated genes involved with cell cycle progression and proliferation, consistent with expression differences in sRCC tumors observed in prior studies^6,10^. The downregulation of the HIF2α gene *EPAS1* with forced expression of *FOSL1* suggests a mechanism by which *FOSL1* may suppress hypoxia-related or angiogenic pathways. This may explain the decreased sensitivity of sRCC to TKIs that target angiogenic pathways and suggests HIF2α inhibition may have less effective in sRCC tumors.

The TFs we implicated in sarcomatoid differentiation through epigenomic profiling showed elevated expression in sRCC tumors across multiple clinical trial cohorts. Patients with high *FOSL1* gene expression levels had better outcomes with ICI+TKIs vs. TKIs alone, whereas those with low *FOSL1* levels had similar outcomes across both treatment arms. This was mainly explained by the decreased benefit of TKIs in *FOSL1*-high patients, mirroring findings in sRCC tumors. Importantly, *FOSL1*-high tumors without sarcomatoid differentiation also demonstrated poor prognosis compared to *FOSL1*-low tumors. This finding raises the possibility that these tumors had epigenomic features of sRCC even though histologic sarcomatoid differentiation was not observed. Our results suggest that clinical stratification based on the expression of sRCC TFs may augment classification provided by histopathologic findings alone.

Because sarcomatoid differentiation is spatially heterogeneous, sampling error poses a significant challenge in the diagnosis of sRCC. Prior studies demonstrated low sensitivity (∼8%)^18^ for detecting sarcomatoid differentiation from tissue biopsies. Accurate diagnosis of sRCC is essential since patients benefit more from ICI-based regimens than TKIs. Since liquid biopsies can sample heterogeneity across tumor sites in advanced cancers^48–51^, a ctDNA-based correlate of sarcomatoid differentiation could overcome issues of sampling error. Utilizing epigenomic profiling of circulating DNA methylation and histone modifications, we detected signatures of sarcomatoid differentiation in patient plasma that distinguished sRCC from epithelioid RCC in plasma samples with low ctDNA (<0.06). Importantly, these signatures were defined entirely using tumor tissue, bolstering biological plausibility and interpretability. In addition, we observed signals of decreased enhancer activity at HIF2α binding sites in sRCC plasma compared to epitheliod RCC plasma, consistent with observations from tumor tissue that HIF2α is downregulated in sRCC. These findings suggest that epigenomic liquid biopsies in RCC could identify patients with sarcomatoid epigenomic features who would benefit from ICI.

Our work has several limitations. Our functional experiments evaluated one TF in a single epithelioid ccRCC cell line (Caki-1). Future work validating other candidate TFs across multiple RCC cell lines is warranted. In addition, our epigenomic profiling from plasma was limited to a small number of plasma samples. Future studies of larger cohorts of plasma samples from patients with sRCC are underway to assess the clinical validity and utility of these findings.

In conclusion, our work demonstrates that sarcomatoid differentiation of RCC is associated with consistent reprogramming of regulatory element activity likely driven by FOSL1 and other EMT-related TFs. We implicated FOSL1 as a predictive biomarker of improved outcomes with ICIs independent of histological evidence of sarcomatoid differentiation. These findings may improve clinical stratification and therapy selection for patients with RCC and inform the discovery of new therapies for sRCC as targeting cancer-associated transcriptional regulation becomes increasingly tractable^52,53^. Finally, our approach represents a broadly applicable framework for using epigenomic liquid biopsy to infer histologic features that can only be assessed from tissue.

## Methods

### Subjects and samples

Sarcomatoid and epithelioid ccRCC clinically annotated tumor tissue specimens were derived as previously described^54^. All tissue samples included were derived from resected RCC tumors from patient donors who provided explicit written consent per the declaration of Helsinki under an approved IRB protocol at the Dana-Farber Cancer Institute. Plasma samples were collected from patients with sarcomatoid and epithelioid ccRCC. The patients were diagnosed and treated at the Dana-Farber Cancer Institute (DFCI) between 2005 and 2022. All patients provided written informed consent. The use of samples was approved by the DFCI (01-045 and 09-171) IRB protocols. Studies were conducted in accordance with recognized ethical guidelines.

### Epigenomic profiling

#### Chromatin immunoprecipitation (ChIP) in RCC tissue specimens

For tissue specimens, a 2-mm^2^ core needle was used to obtain one core of tumor tissue from frozen tissue blocks in the areas marked on the corresponding slide enriched for tumor cells of sarcomatoid and non-sarcomatoid regions. Frozen samples were pulverized using the Covaris CryoPrep system. They were then fixed using 2 mmol/L disuccinimidyl glutarate (DSG) for 10 minutes, followed by 1% formaldehyde buffer for 10 minutes, and quenched with glycine. Chromatin was sheared to 300 to 500 bp using the Covaris E220 ultrasonicator. The resulting chromatin was incubated overnight with the following antibodies (H3K27ac, Diagenode, Catalog No: C15410196 LOT: A1723-0041D; H3K4me2, Diagenode, Catalog No: C15410035 LOT: A936-0023) coupled with 40 μl protein A and protein G beads (Invitrogen) at 4 degrees Celsius overnight. Five percent of the sample was not exposed to antibodies and was used as a control input. The beads were then washed three times each with Low-Salt Wash Buffer (0.1% SDS, 1% Triton X-100, 2 mM EDTA, 20 mM Tris-HCl pH 7.5, 150 mM NaCl), High-Salt Wash Buffer (0.1% SDS, 1% Triton X-100, 2 mM EDTA, 20 mM Tris-HCl pH 7.5, 500 mM NaCl), and LiCl Wash Buffer (10 mM Tris pH 7.5, 250 mM LiCl, 1% NP-40, 1% Na-Doc, 1 mM EDTA) and rinsed with TE buffer (pH 8.0) once. The samples were then de-cross-linked, treated with RNase and proteinase K, and DNA was extracted (Qiagen). DNA sequencing libraries were prepared from purified input and IP sample DNA using the ThruPLEX DNA-seq Kit (TakaraBio). Libraries were sequenced on an Illumina HiSeq 4000 to generate 150-bp paired-end reads (Novogene).

#### ChIP-seq data analysis

ChIP-sequencing reads were aligned to the human genome build hg19 using the Burrows-Wheeler Aligner (BWA) version 0.7.17^55^. Non-uniquely mapped and redundant reads were discarded. MACS v2.1.1.20140616^56^ was used for ChIP-seq peak calling with a q-value (FDR) threshold of 0.01. ChIP-seq data quality was evaluated by a variety of measures, including total peak number, FrIP (fraction of reads in peak) score, number of high-confidence peaks (enriched >10-fold over background), and percent of peak overlap with DNAse hypersensitivity (DHS) peaks derived from the ENCODE project. ChIP-seq peaks were assessed for overlap with gene features and CpG islands using annotatr^57^. IGV v2.8.2^58^ was used to visualize normalized ChIP-seq read counts at specific genomic loci. ChIP-seq heatmaps were generated with deepTools v3.3.1^59^ and show normalized read counts at the peak center ±2 kb unless otherwise noted. Overlap of ChIP-seq peaks was assessed using BEDTools v2.26.0. Peaks were considered overlapping if they shared one or more base pairs.

#### Identification and annotation of histology-specific peaks

Sample–sample clustering, principal component analysis, and identification of lineage-enriched peaks were performed using Cobra v2.0^60^ (https://bitbucket.org/cfce/cobra/src/master/), a ChIP-seq analysis pipeline implemented with Snakemake^61^. ChIP-seq data from sarcomatoid and epithelioid RCC tissue samples were compared to identify H3K27ac and H3K4me2 peaks with significant enrichment in the two above groups. Samples from unique individuals were included. A union set of peaks for each histone modification was created using BEDTools, and narrowPeak calls from MACS were used for H3K27ac and H3K4me2. The number of unique aligned reads overlapping each peak in each sample was calculated from BAM files using BEDtools. Read counts for each peak were normalized to the total number of mapped reads for each sample. Quantile normalization was applied to this matrix of normalized read counts. Using DEseq2 v1.14.1^62^, histology-enriched peaks were identified at the indicated FDR-adjusted *p*-value (*p*_adj_) and log2 fold-change cutoffs (H3K27ac, *p*_adj_ < 0.05, |log2 fold-change| >0; H3K4me2, *p*_adj_ < 0.05, |log2 fold-change| >0;). Unsupervised hierarchical clustering was performed based on Spearman correlation between samples. Principal component analysis was performed using the prcomp R function. Enriched de novo motifs in differential peaks were detected using HOMER version 4.7. The top non-redundant motifs were ranked by adjusted *p*-value.

The GREAT analysis^31^ (V3.0) was used to assess for enrichment of Gene Ontology (GO) and MSigDB perturbation annotations among genes near differential ChIP-seq peaks, assigning each peak to the nearest gene within 500 kb.

#### TCGA and Clinical Trials Data

RNA-seq and clinical data from Javelin Renal101, IMmotion151 and TCGA cohorts were analyzed using R (v 4.2) on Rstudio (v 2022.7.2.576). Differential gene expression analysis between sRCC and epithelioid ccRCC tissue samples was computed using DESeq2 and Benjamini-Hochberg false discovery rate correction with q < 0.05 considered statistically significant^62^. Patients were then stratified based on transcript per millions (TPMs) counts for transcription factors of interest. Survival analysis was computed using survminer R package with p < 0.05 considered statistically significant.

#### Generation of stable CRISPRa FOSL1 cell lines

Guides were cloned into a pXPR_502, following a previously published cloning protocol^63^: sgCtrl: F: 5’-CACCGCGCCAAACGTGCCCTGACGG-3’, R: 5’-AAACCCGTCAGGGCACGTTTGGCGC-3’, sgFOSL1: F: 5’-CACCGGGGCTGAACCACTGCGACCG-3’, R: 5’-AAACCGGTCGCAGTGGTTCAGCCCC-3’ 24 h before transfection, HEK293T cells were seeded 10cm dishes at a density of 5 × 10^6^ cells. PEI-Transfection was performed following the manufacturer’s protocol. Briefly, one solution of Opti-MEM (100 μL) and PEI MAX (40μL) was combined with a DNA mixture of the packaging plasmid pMD2.G (2μg), psPAX2 (3 μg), and the transfer vector (5 μg). This mixture was incubated at room temperature 15 minutes and added dropwise on HEK293T cells with fresh media. After an overnight incubation at 37°C, the media was changed and collected after 48 hours. The virus was then concentrated in Amicon® Ultra-15 50 kDa, at 1500g for 30 min and stored at −80°C.

Lentiviral infection was performed on Caki-1 cells with the following conditions: 200μL of concentrated virus was added to 2 x 10^5^ cells in replicates with 4μg/mL of polybren in 6 well-dishes with 2 mL of media, centrifugated at 1000g during 1h. The media was changed after an overnight incubation, and antibiotic selection started after 48 hours, for 5 days.

Cakif-1 were first infected with pXPR_109 (dCas9-VP64) and selected with blasticidin (5μg/mL) to establish stable dCas9-VP64-expressing Caki-1 cells. Subsequently, these cells were infected with pXPR-502 containing guides and selected with puromycin (2 μg/mL) and blasticidin to maintain the dCas9-VP64 expression.

#### RNA methods

RNA was extracted using the Qiagen RNeasy Mini Kit (Cat No./ID: 74104) as recommended, from frozen tumor samples adjacent to those samples used for ChIP. RNA-seq libraries were constructed from 1 μg total RNA using the Illumina TruSeq Stranded mRNA LT Sample Prep Kit according to the manufacturer’s protocol. Barcoded libraries were pooled and sequenced on the Illumina HiSeq 2500 generating 50-bp paired-end reads.

#### Functional validation RNA-seq data

Differential gene expression analysis between FOSL1 CRISPRa and negative control Caki-1 cells was computed using DESeq2 and Benjamini-Hochberg false discovery rate correction with q < 0.05 considered statistically significant^62^. Gene Set Enrichment Analysis (GSEA) was computed between FOSL1 CRISPRa and negative control Caki-1 cells (GSEA q < 0.01) as previously described^64^.

#### cfDNA processing and tumor content calculation

cfDNA samples were processed by the following method. Peripheral blood was collected in EDTA Vacutainer tubes (BD) and processed within 3 hours of collection. Plasma was separated by centrifugation at 2,500 × *g* for 10 minutes, transferred to microcentrifuge tubes, and centrifuged at 2,500 × *g* at room temperature for 10 minutes to remove cellular debris. The supernatant was aliquoted into 1 to 2 mL aliquots and stored at −80°C until DNA extraction. cfDNA was isolated from 1 mL of plasma using the QIAGEN Circulating Nucleic Acids Kit (QIAGEN), eluted in AE buffer, and stored at −80°C. Low-pass whole-genome sequencing (LPWGS) was performed on all cfDNA samples. The ichorCNA R package was used to infer copy-number profiles and cfDNA tumor content from read abundance across bins spanning the genome using default parameters^65^.

#### MeDIP-seq

MeDIP-seq was performed on tissue and plasma samples using previously published methods^66^. cfDNA library preparation was performed on 10 ng of DNA using the KAPA HyperPrep Kit (KAPA Biosystems) according to the manufacturer’s protocol. We then performed end-repair, A-tailing, and ligation of NEBNext adaptors (NEBNext Multiplex Oligos for Illumina kit, New England BioLabs). Libraries were digested using the USER enzyme (New England BioLabs). λ DNA, consisting of unmethylated and *in vitro* methylated DNA, was added to prepared libraries to achieve a total amount of 100 ng DNA. Methylated and unmethylated *Arabidopsis thaliana* DNA (Diagenode) was added for quality control. MeDIP was performed using the MagMeDIP Kit (Diagenode) following the manufacturer’s protocol. Samples were purified using the iPure Kit v2 (Diagenode). Success of the immunoprecipitation was confirmed using qPCR to detect recovery of the spiked-in *Arabidopsis thaliana* methylated and unmethylated DNA. KAPA HiFi Hotstart ReadyMix (KAPA Biosystems) and NEBNext Multiplex Oligos for Illumina (New England Biolabs) were added to a final concentration of 0.3 μmol/L. Libraries were amplified as follows: activation at 95°C for 3 minutes, amplification cycles of 98°C for 20 seconds, 65°C for 15 seconds, 72°C for 30 seconds, and a final extension of 72°C for 1 minute. Samples were pooled and sequenced (Novogene Corporation) on Illumina HiSeq 4000 to generate 150 bp paired-end reads.

#### MeDIP-seq quality control and processing of sequencing reads

After sequencing, the quality and quantity of the raw reads were examined using FastQC version 0.11.5 (http://www.bioinformatics.babraham.ac.uk/projects/fastqc) and MultiQC version 1.7^67^. Raw reads were quality, and adapter trimmed using Trim Galore! version 0.6.0 (http://www.bioinformatics.babraham.ac.uk/projects/trim_galore/) using default settings in paired-end mode. The trimmed reads then were aligned to hg19 using Bowtie2 version 2.3.5.1 in paired-end mode and all other settings default^68^. The SAMtools version 1.10 software suite was used to convert SAM alignment files to BAM format, sort and index reads, and remove duplicates^69^. The R package RSamtools version 2.2.1 was used to calculate the number of unique mapped reads. Saturation analyses to evaluate reproducibility of each library were carried out using the R Bioconductor package MEDIPS version 1.38.0^70^.

#### Tissue-informed approach for detection of sarcomatoid features using MeDIP-seq

To identify differentially methylated regions (DMR) between sarcomatoid and non-sarcomatoid ccRCC samples, we first binned the genome into 300 base-pair windows and tested each window for differential methylation between sarcomatoid and non-sarcomatoid samples using limma-voom (R package limma version 3.42.0) on TMM-normalized counts (R package edgeR version 3.28.0)^71,72^. Only bins with a total count above a fixed threshold were tested for differential methylation, where the threshold was set at 20% of the total number of samples across both groups. We restricted our search to bins within annotated CpG islands and FANTOM5 enhancers and excluded regions of high signal or poor mappability ^57,73^. We selected DMRs with read enrichment in sRCC compared with epithelioid RCC at FDR-adjusted *P* < 0.01 and log_2_ fold-change > 0. We removed windows with peaks in MeDIP-seq data from white blood cells (as determined by MACS2, version 2.1.2) to minimize signal from blood cell–derived cfDNA^56^. Using the MeDIPs R package, we calculated CpG-normalized relative methylation scores (RMS) across 300 bp windows for each cfDNA sample^70,74^. We then summed RMS in cfDNA at sarcomatoid-enriched DMRs for each sample and normalized this value to the sum of RMS values across all 300 bp windows. This value was termed “Sarcomatoid RCC Methylation Value.” The same process was performed for Non-sarcomatoid RCC DMRs to derive a “Non-sarcomatoid RCC Methylation Value.” We then calculated the log_2_ ratio of the Sarcomatoid RCC Methylation Value to the Non-sarcomatoid RCC Methylation Value and normalized these values to the median score in cfDNA from eight healthy cancer-free controls. This value was termed the “Sarcomatoid RCC Risk Score.”

##### cfChIP-seq assay

1 μg of antibody was coupled with 10 μL protein A (Invitrogen, cat# 10002D) and 10 μL protein G (Invitrogen, cat# 10004D) for at least 6 hrs at 4 °C with rotation in 0.5 % BSA (Jackson Immunology, cat# 001-000-161) in PBS (Gibco, cat# 14190250), followed by blocking with 1% BSA in PBS for 1 hr at 4°C with rotation. The following antibodies were used: H3K4me3, Thermo Fisher # PA5-27029; H3K27ac, Abcam # ab4729; panAc, Active Motif #39139. Thawed plasma was centrifuged at 3,000g for 15 min at 4°C. The supernatant was precleared with the magnetic beads with 20 μL protein A and 20 μL protein G for 2 hrs at 4°C. Then, the precleared and conditioned plasma was subjected to antibody-coupled magnetic beads overnight with rotation at 4 °C. The reclaimed magnetic beads were washed with 1mL of each washing buffer twice. Three washing buffers were used in following order: low salt washing buffer (0.1% SDS, 1% Triton X-100, 2 mM EDTA, 150mM NaCl, 20 mM Tris-HCl pH 7.5), high salt buffer (0.1 % SDS, 1 % Triton X-100, 2 mM EDTA, 500 mM NaCl, 20 mM Tris-HCl pH 7.5), and LiCl washing buffer (250 mM LiCl, 1%NP-40, 1% Na Deoxycholate, 1 mM EDTA, 10 mM Tris-HCl pH 7.5). Subsequently, the beads were rinsed with TE buffer (Fisher Sci, cat# BP2473500), and resuspended and incubated in 100μL of DNA extraction buffer containing 0.1 M NaHCO3, 1% SDS and 0.6 mg/mL Proteinase K (Qiagen, cat#19131) and 0.4 mg/mL RNaseA (Thermo Fisher, cat#12091021) for 10 min for 37°C, for 1 hr for 50°C, and for 90 min at 65°C. DNA was purified through Phenol extraction (Invitrogen, cat# 15593031) and Ethanol precipitation was performed with 3M NaOAc (Ambion, cat# AM9740) and glycogen (Ambion, cat# AM9510). cfChIP-seq libraries were prepared with ThruPLEX DNA-Seq Kit (Takara Bio, cat# R400675) following the manufacturer’s instructions. After library amplification, the DNA was purified by AMPure XP (Beckman coulter, cat# A63880). The size distribution of the purified libraries was examined using Agilent 2100 Bioanalyzer with a high sensitivity DNA Chip (Agilent, cat# 5067-4626). The library was submitted for the 150 base-pair paired end sequencing on an Illumina NovaSeq6000 system (Novogene Corporation, CA).

### Statistical tests

All statistical tests were two-sided except where otherwise indicated.

## Supporting information

Supplementary Data S1. Clinical and pathological data of tissue samples undergoing epigenomic profiling.

Supplementary Data S2. Association of expression levels of transcription factors with clinical outcomes.

Supplementary Data S3. Association of FOSL1 expression with clinical outcomes by treatment arm.

Supplementary Data S4. Clinical and pathologic features of plasma samples undergoing epigenomic profiling.

## Supplementary Material

### Supplementary Figures

**Supplementary Figure 1. Epithelioid and sarcomatoid ccRCC tissue samples have distinct epigenomic profiles. a-c** Unsupervised hierarchical clustering of the H3K27ac, H3K4me2, and MeDIP peaks in epithelioid and sarcomatoid ccRCC tissue samples.

**Supplementary Figure 2. Clinical outcomes of patients with RCC by *FOSL1* levels. a** Progression-free survival in patients in IM151 by *FOSL1* levels. **b** Progression-free survival in patients with sRCC and epithelioid ccRCC divided by expression of *FOSL1* (High vs. Low) in the avelumab plus axitinib and atezolizumab plus bevacizumab arms of JR101 and IM151, respectively. TPM: Transcripts per million, TCGA: The Cancer Genome Atlas, JR101: Javelin Renal 101, IM151: IMmotion151, AveAxi: avelumab plus axitinib. AtezoBev: atezolizumab plus bevacizumab.

**Supplementary Figure 3. Impact of activation of *FOSL1* gene expression on epigenomic and transcriptomic profiles of Caki-1 cells. a** H3K27ac signal at promoter sites of genes upregulated in *FOSL1* CRISPRa and control Caki-1 cells. **b** Gene set enrichment analysis (GSEA) of genes upregulated in sarcomatoid ccRCC from JR101 in *FOSL1* CRISPRa Caki-1 cells.

### Supplementary Data

**Supplementary Data S1.** Clinical and pathological data of tissue samples undergoing epigenomic profiling.

**Supplementary Data S2.** Association of expression levels of transcription factors with clinical outcomes.

**Supplementary Data S3.** Association of *FOSL1* expression with clinical outcomes by treatment arm.

**Supplementary Data S4.** Clinical and pathologic features of plasma samples undergoing epigenomic profiling.

## Notes

### Competing Interest Statement

S.R.V. acknowledges Support from the Doris Duke Charitable Foundation (Clinical Scientist Development Award, 2020101), R01CA279044, and the V Foundation (V Scholar Award, V2022-018). S.R.V.: Consulting (current or past 3 years): Jnana Therapeutics. Research support from Bayer. Spouse is an employee of and holds equity in Kojin Therapeutics. D.A.B. reports honoraria from LM Education/Exchange Services, advisory board fees from Exelixis and AVEO, consulting fees from Merck, Pfizer, and Elephas, equity in Elephas, Fortress Biotech (subsidiary), and CurIOS Therapeutics, personal fees from Schlesinger Associates, Cancer Expert Now, Adnovate Strategies, MDedge, CancerNetwork, Catenion, OncLive, Cello Health BioConsulting, PWW Consulting, Haymarket Medical Network, Aptitude Health, ASCO Post/Harborside, Targeted Oncology, Accolade 2nd.MD, DLA Piper, AbbVie, Compugen, Link Cell Therapies, and Scholar Rock, and research support from Exelixis and AstraZeneca, outside of the submitted work. S.C.B., T.K.C. and M.L.F. are co-founders and shareholders of Precede Biosciences. SCB is supported by grant W81XWH-21-1-0299 from the Department of Defense, a Clinical Investigator Award from the Damon Runyon Cancer Research Foundation, the Kure It Cancer Research Foundation, and the Fund for Innovation in Cancer Informatics. The remaining authors report no competing interests.

