## Supplementary figures and images for "Epigenomic signatures as circulating and predictive biomarkers in sarcomatoid renal cell carcinoma"

### Supplementary Data S1. Clinical and pathological data of tissue samples undergoing epigenomic profiling.

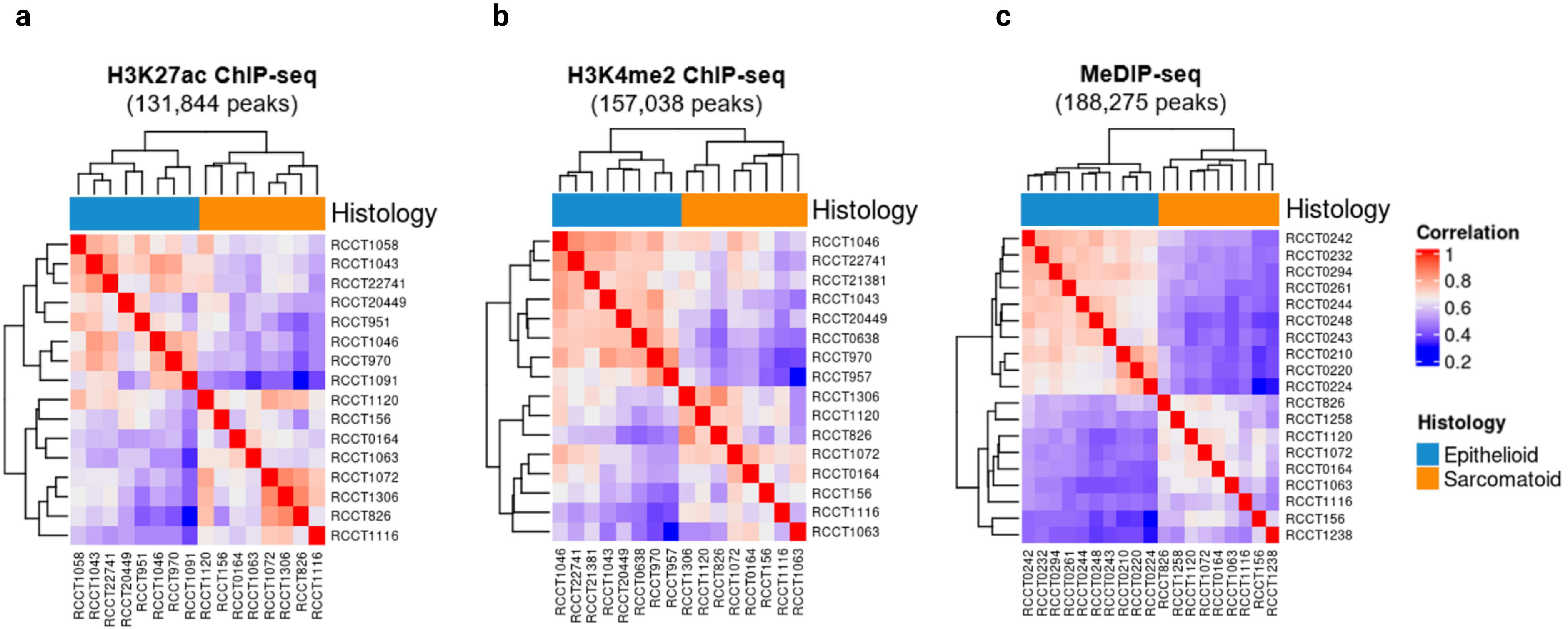

### Supplementary Data S2. Association of expression levels of transcription factors with clinical outcomes.

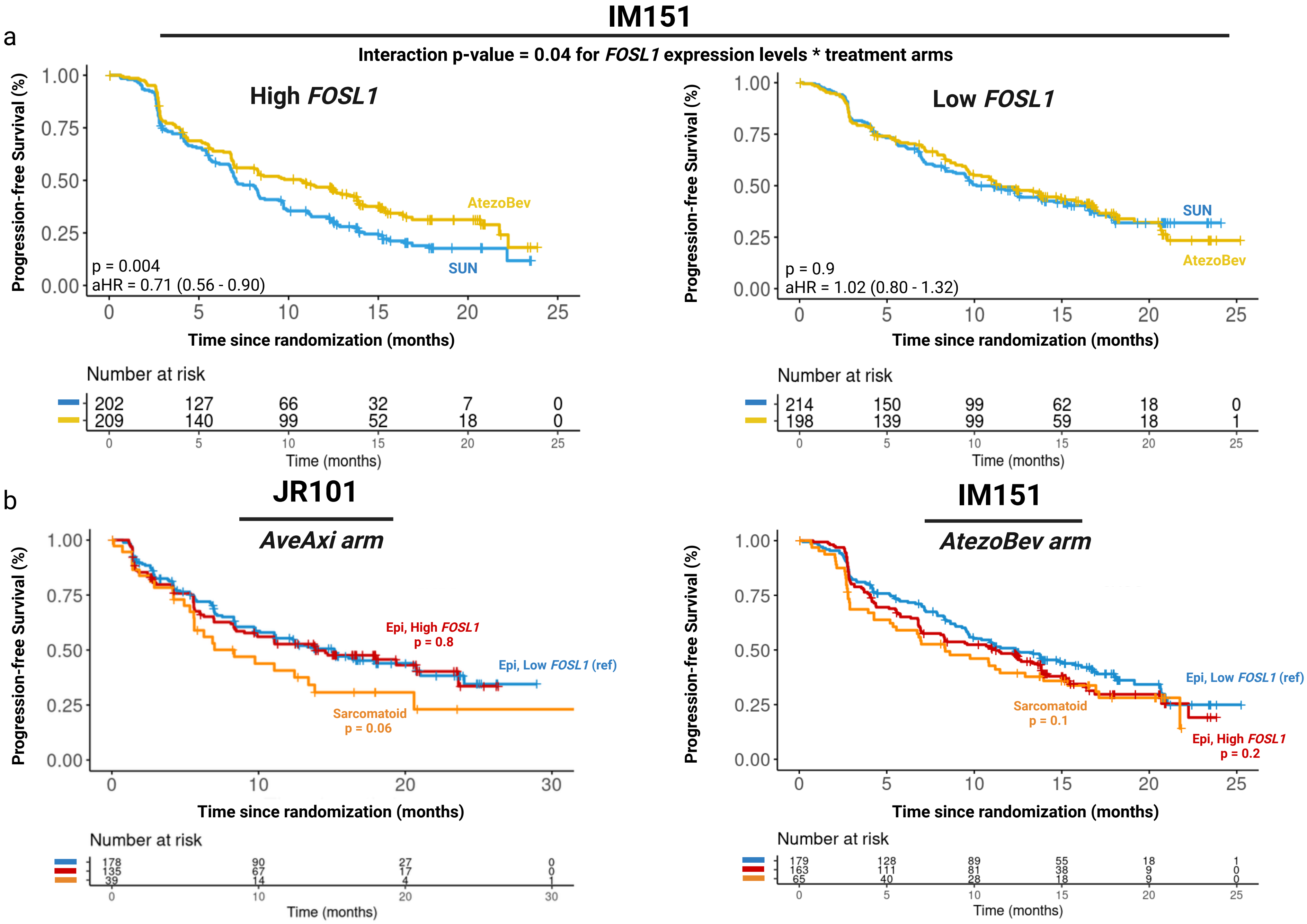

### Supplementary Data S3. Association of FOSL1 expression with clinical outcomes by treatment arm.

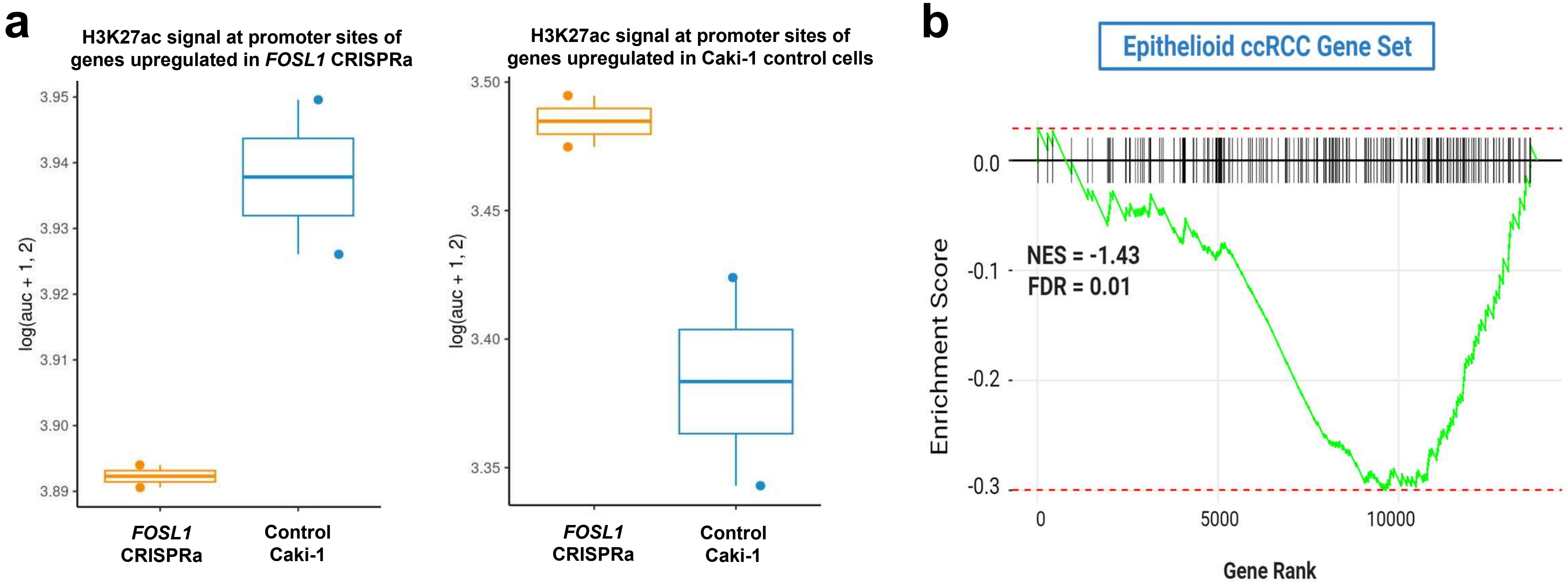
