## Supplementary Data S4. Clinical and pathologic features of plasma samples undergoing epigenomic profiling. for "Epigenomic signatures as circulating and predictive biomarkers in sarcomatoid renal cell carcinoma"

**Supplementary Data S1.** Clinical and pathological data of tissue samples undergoing epigenomic profiling.

| Sample | Sex | Age | Stage | Grade | Sarcomatoid Differentiation (%) | Tissue Source | H3K27Ac | H3K4me2 | MeDIP |
|---|---|---|---|---|---|---|---|---|---|
| RCCT1189 | Female | 50-59 | T3aN0 | III | No | Metastatic | Yes | No | No |
| RCCT951 | Male | 50-59 | T3aN0 | II | No | Primary | Yes | No | No |
| RCCT0638 | Male | 60-69 | T3Nx | II | No | Primary | No | Yes | No |
| RCT1320 | Female | 60-69 | T3aNx | III | No | Metastatic | Yes | No | No |
| RCCT21381 | Male | 40-49 | T3bN1 | II | No | Primary | No | Yes | No |
| RCCT957 | Male | 40-49 | T3aNx | III | No | Primary | No | Yes | No |
| RCCT970 | Female | 50-59 | T3aN0M1 | III | No | Primary | Yes | Yes | No |
| RCCT22741 | Female | 50-59 | T3aNx | IV | No | Primary | Yes | Yes | No |
| RCCT1046 | Male | 60-69 | T3cNx | III | No | Primary | Yes | Yes | No |
| RCCT20449 | Male | 60-69 | T3a | III | No | Primary | Yes | Yes | No |
| RCCT1043 | Female | 70-79 | T3aNx | IV | No | Primary | Yes | Yes | No |
| RCCT0826 | Male | 70-79 | T3aNx | IV | Yes (focal) | Primary | Yes | Yes | Yes |
| RCCT0156 | Male | 60-69 | T2N0Mx | IV | Yes (focal) | Primary | Yes | Yes | Yes |
| RCCT0164 | Female | 60-69 | T2N2Mx | IV | Yes (extensive) | Primary | Yes | Yes | Yes |
| RCCT1063 | Female | 50-59 | T4N1M0 | IV | Yes (focal) | Primary | Yes | Yes | Yes |
| RCCT1072 | Male | 60-69 | T3aNx | IV | Yes (<5%) | Primary | Yes | Yes | Yes |
| RCCT1116 | Male | 50-59 | T3aN0 | IV | Yes (<10%) | Primary | Yes | Yes | Yes |
| RCCT1238 | Male | 50-59 | T4NxM1 | IV | Yes (25%) | Metastatic | No | No | Yes |
| RCCT1258 | Male | 70-79 | T3aN1M0 | IV | Yes (70%) | Primary | No | No | Yes |
| RCCT1306 | Male | 30-39 | T3aNx | IV | Yes (40%) | Primary | Yes | Yes | No |
| RCCT1120 | Male | 50-59 | T2aN0M1 | IV | Yes (focal) | Metastatic | Yes | Yes | Yes |
| RCCT0210 | Male | 60-69 | T1aNxMX | III | No | Primary | No | No | Yes |
| RCCT0220 | Male | 80-89 | T1aNxMX | II | No | Primary | No | No | Yes |
| RCCT0224 | Male | 40-50 | T1aNxMX | II | No | Primary | No | No | Yes |
| RCCT0232 | Female | 60-69 | T1bN0MX | II | No | Primary | No | No | Yes |
| RCCT0242 | Male | 60-69 | T1aNXMX | III | No | Primary | No | No | Yes |
| RCCT0243 | Female | 50-59 | T1N0MX | II | No | Primary | No | No | Yes |
| RCCT0244 | Female | 70-79 | T1N0MX | II | No | Primary | No | No | Yes |
| RCCT0248 | Male | 40-49 | T1aNXMX | II | No | Primary | No | No | Yes |
| RCCT0261 | Male | 60-69 | T1aNxMX | II | No | Primary | No | No | Yes |
| RCCT0294 | Male | 60-69 | T1aNxMX | II | No | Primary | No | No | Yes |

**Supplementary Data S2.** Association of expression levels of transcription factors with clinical outcomes.

| | JR101 (AveAxi vs. Sunitinib) | | | | | IM151 (AtezoBev vs. Sunitinib) | | | | |
|---|---|---|---|---|---|---|---|---|---|---|
| | High | | Low | | | High | | Low | | |
| **TF** | **HR*** | **p-value** | **HR*** | **p-value** | **interaction p-value** | **HR*** | **p-value** | **HR*** | **p-value** | **interaction p-value** |
| FOSL1 | 0.53 (0.41 - 0.70) | **0.000009** | 0.85 (0.63 - 1.13) | 0.3 | **0.03** | **0.71 (0.56 - 0.90)** | **0.004** | 1.02 (0.80 - 1.32) | 0.9 | **0.04** |
| E2F7 | 0.61 (0.47 - 0.80) | **0.0003** | 0.78 (0.58 - 1.04) | 0.09 | 0.18 | **0.76 (0.60 - 0.96)** | **0.02** | 1.00 (0.77 - 1.28) | 0.97 | 0.08 |
| ETV4 | 0.58 (0.44 - 0.77) | **0.0002** | 0.77 (0.58 - 1.01) | 0.1 | 0.14 | **0.85 (0.67 - 1.07)** | 0.16 | 0.82 (0.63 - 1.06) | 0.13 | 0.9 |

*HR TKI_ICI vs. TKI alone for PFS adjusted for IMDC risk groups

**Supplementary Data S3.** Association of *FOSL1* expression with clinical outcomes by treatment arm.

**JR101**

| TF | Treatment Arm | Group | N | Median PFS, Months | HR_PFS(95%CI)* | p-value |
|---|---|---|---|---|---|---|
| FOSL1 | AveAxi | LowTPM non-sRCC | 178 | 15.2 | Reference | |
| | | HighTPM non-sRCC | 135 | 13.9 | 0.91(0.66-1.25) | 0.55 |
| | | sRCC | 39 | 8.3 | 1.31(0.84-2.05) | 0.24 |
| | SUN | LowTPM non-sRCC | 160 | 11.3 | Reference | |
| | | HighTPM non-sRCC | 155 | 6.5 | 1.54(1.14-2.06) | **0.004** |
| | | sRCC | 56 | 4.1 | 2.01(1.36-2.97) | **<0.001** |

**IM151**

| TF | Treatment Arm | Group | N | Median PFS, Months | HR_PFS(95%CI)* | p-value |
|---|---|---|---|---|---|---|
| FOSL1 | AtezoBev | LowTPM non-sRCC | 179 | 12.4 | Reference | |
| | | HighTPM non-sRCC | 163 | 11.2 | 1.11(0.85-1.47) | 0.44 |
| | | sRCC | 65 | 8.3 | 1.02(0.70-1.47) | 0.94 |
| | SUN | LowTPM non-sRCC | 197 | 11.8 | Reference | |
| | | HighTPM non-sRCC | 149 | 8.3 | 1.35(1.03-1.76) | **0.030** |
| | | sRCC | 69 | 5.6 | 2.03(1.47-2.79) | **<0.001** |

*Adjusted for IMDC risk groups

**Supplementary Data S4.** Clinical and pathologic features of plasma samples undergoing epigenomic profiling.

| Study_ID | Source | RCC vs. Healthy | Histology | Sarcomatoid Features | Antibody | Peak Number | ctDNA | Enrichment | Fragments |
|---|---|---|---|---|---|---|---|---|---|
| K27_RCC1764 | DFCI | RCC | Clear cell | Yes | H3K27ac | 6573 | 0.00 | 12.3 | 446055 |
| K27_RCC2031 | DFCI | RCC | Clear cell | Yes | H3K27ac | 1685 | 0.06 | 17.2 | 2092763 |
| K27_RCC2194 | DFCI | RCC | Clear cell | Yes | H3K27ac | 766 | 0.00 | 6.7 | 4072253 |
| K27_RCC478 | DFCI | RCC | Clear cell | Yes | H3K27ac | 2641 | 0.00 | 6.6 | 846586 |
| K27_RCC594 | DFCI | RCC | Clear cell | Yes | H3K27ac | 1980 | 0.04 | 8.3 | 1771094 |
| K27_RCC2970 | DFCI | RCC | Clear cell | No | H3K27ac | 4862 | 0.03 | 23.7 | 8584283 |
| K27_RCC2973 | DFCI | RCC | Clear cell | No | H3K27ac | 712 | 0.03 | 7.1 | 10347375 |
| K27_RCC2971 | DFCI | RCC | Clear cell | No | H3K27ac | 613 | 0.00 | 4.9 | 6045526 |
| K27_RCC2974 | DFCI | RCC | Clear cell | No | H3K27ac | 2850 | 0.05 | 13.4 | 12426839 |
| K27_RCC2975 | DFCI | RCC | Clear cell | No | H3K27ac | 3655 | 0.04 | 10.0 | 19571161 |
| K27_HP030132 | MGB | Healthy | N/A | N/A | H3K27ac | 9112 | 0.00 | 42.8 | 3569398 |
| K27_HP030642 | MGB | Healthy | N/A | N/A | H3K27ac | 9179 | 0.00 | 41.3 | 3653087 |
| K27_HP031645 | MGB | Healthy | N/A | N/A | H3K27ac | 14221 | 0.00 | 13.7 | 10796830 |
| K27_HP034881 | MGB | Healthy | N/A | N/A | H3K27ac | 10811 | 0.00 | 21.9 | 6784781 |
| K27_HP035094 | MGB | Healthy | N/A | N/A | H3K27ac | 13998 | 0.00 | 22.5 | 7410494 |
| K27_HP038748 | MGB | Healthy | N/A | N/A | H3K27ac | 8754 | 0.00 | 40.9 | 6036870 |
| K27_HP041556 | MGB | Healthy | N/A | N/A | H3K27ac | 10727 | 0.00 | 5.9 | 17101237 |
| K27_HP056703 | MGB | Healthy | N/A | N/A | H3K27ac | 10848 | 0.00 | 26.6 | 7628759 |
| K27_HP098228 | MGB | Healthy | N/A | N/A | H3K27ac | 12748 | 0.00 | 24.8 | 6281549 |
| K4_RCC1764 | DFCI | RCC | Clear cell | Yes | H3K4me3 | 7613 | 0.00 | 12.3 | 385108 |
| K4_RCC2031 | DFCI | RCC | Clear cell | Yes | H3K4me3 | 11591 | 0.06 | 16.9 | 4800904 |
| K4_RCC2194 | DFCI | RCC | Clear cell | Yes | H3K4me3 | 1431 | 0.00 | 8.1 | 1907827 |
| K4_RCC478 | DFCI | RCC | Clear cell | Yes | H3K4me3 | 2486 | 0.00 | 4.4 | 1219084 |
| K4_RCC594 | DFCI | RCC | Clear cell | Yes | H3K4me3 | 576 | 0.04 | 4.4 | 6185859 |
| K4_RCC2970 | DFCI | RCC | Clear cell | No | H3K4me3 | 11505 | 0.03 | 22.2 | 2571935 |
| K4_RCC2973 | DFCI | RCC | Clear cell | No | H3K4me3 | 1432 | 0.03 | 4.1 | 9389360 |
| K4_RCC2971 | DFCI | RCC | Clear cell | No | H3K4me3 | 2076 | 0.00 | 10.0 | 2108453 |
| K4_RCC2974 | DFCI | RCC | Clear cell | No | H3K4me3 | 9004 | 0.05 | 20.0 | 3025504 |
| K4_RCC2975 | DFCI | RCC | Clear cell | No | H3K4me3 | 8903 | 0.04 | 16.3 | 2253785 |
| K4_HP030132 | MGB | Healthy | N/A | N/A | H3K4me3 | 15751 | 0.00 | 30.0 | 4854960 |
| K4_HP030642 | MGB | Healthy | N/A | N/A | H3K4me3 | 16262 | 0.00 | 48.7 | 3442523 |
| K4_HP031645 | MGB | Healthy | N/A | N/A | H3K4me3 | 18649 | 0.00 | 33.9 | 8834730 |
| K4_HP034881 | MGB | Healthy | N/A | N/A | H3K4me3 | 17574 | 0.00 | 40.8 | 5534610 |
| K4_HP035094 | MGB | Healthy | N/A | N/A | H3K4me3 | 15478 | 0.00 | 41.2 | 4243016 |
| K4_HP038748 | MGB | Healthy | N/A | N/A | H3K4me3 | 20013 | 0.00 | 42.6 | 6359502 |
| K4_HP041556 | MGB | Healthy | N/A | N/A | H3K4me3 | 16890 | 0.00 | 19.5 | 7176560 |
| K4_HP056703 | MGB | Healthy | N/A | N/A | H3K4me3 | 16210 | 0.00 | 34.0 | 4106019 |
| K4_HP098228 | MGB | Healthy | N/A | N/A | H3K4me3 | 16160 | 0.00 | 35.2 | 4775747 |
| mdRCC1764 | DFCI | RCC | Clear cell | Yes | MeDIP | 18754 | 0.00 | N/A | 4221011 |
| mdRCC2031 | DFCI | RCC | Clear cell | Yes | MeDIP | 152418 | 0.06 | N/A | 5026227 |
| mdRCC2194 | DFCI | RCC | Clear cell | Yes | MeDIP | 70634 | 0.00 | N/A | 8100673 |

| | | | | | | | | | |
|---|---|---|---|---|---|---|---|---|---|
| mdRCC478 | DFCI | RCC | Clear cell | Yes | MeDIP | 17297 | 0.00 | N/A | 4278345 |
| mdRCC594 | DFCI | RCC | Clear cell | Yes | MeDIP | 24194 | 0.04 | N/A | 4780142 |
| mdRCC2970 | DFCI | RCC | Clear cell | No | MeDIP | 76389 | 0.03 | N/A | 6497492 |
| mdRCC2973 | DFCI | RCC | Clear cell | No | MeDIP | 114175 | 0.03 | N/A | 7956452 |
| mdRCC2971 | DFCI | RCC | Clear cell | No | MeDIP | 103694 | 0.00 | N/A | 7450487 |
| mdRCC2974 | DFCI | RCC | Clear cell | No | MeDIP | 47902 | 0.05 | N/A | 4906426 |
| mdRCC2975 | DFCI | RCC | Clear cell | No | MeDIP | 74826 | 0.04 | N/A | 6960489 |
| mdHP030642 | MGB | Healthy | N/A | N/A | MeDIP | 51831 | 0.00 | N/A | 4273099 |
| mdHP031645 | MGB | Healthy | N/A | N/A | MeDIP | 188035 | 0.00 | N/A | 18432797 |
| mdHP034881 | MGB | Healthy | N/A | N/A | MeDIP | 87583 | 0.00 | N/A | 6389812 |
| mdHP035094 | MGB | Healthy | N/A | N/A | MeDIP | 122896 | 0.00 | N/A | 10207526 |
| mdHP038748 | MGB | Healthy | N/A | N/A | MeDIP | 179435 | 0.00 | N/A | 18361190 |
| mdHP041556 | MGB | Healthy | N/A | N/A | MeDIP | 173422 | 0.00 | N/A | 16120905 |
| mdHP056703 | MGB | Healthy | N/A | N/A | MeDIP | 41449 | 0.00 | N/A | 3658589 |
| mdHP098228 | MGB | Healthy | N/A | N/A | MeDIP | 22848 | 0.00 | N/A | 3646858 |
| LRCC1764 | DFCI | RCC | Clear cell | Yes | LPWGS | N/A | 0.00 | N/A | N/A |
| LRCC2031 | DFCI | RCC | Clear cell | Yes | LPWGS | N/A | 0.06 | N/A | N/A |
| LRCC2194 | DFCI | RCC | Clear cell | Yes | LPWGS | N/A | 0.00 | N/A | N/A |
| LRCC478 | DFCI | RCC | Clear cell | Yes | LPWGS | N/A | 0.00 | N/A | N/A |
| LRCC594 | DFCI | RCC | Clear cell | Yes | LPWGS | N/A | 0.04 | N/A | N/A |
| LRCC2970 | DFCI | RCC | Clear cell | No | LPWGS | N/A | 0.03 | N/A | N/A |
| LRCC2973 | DFCI | RCC | Clear cell | No | LPWGS | N/A | 0.03 | N/A | N/A |
| LRCC2971 | DFCI | RCC | Clear cell | No | LPWGS | N/A | 0.00 | N/A | N/A |
| LRCC2974 | DFCI | RCC | Clear cell | No | LPWGS | N/A | 0.05 | N/A | N/A |
| LRCC2975 | DFCI | RCC | Clear cell | No | LPWGS | N/A | 0.04 | N/A | N/A |
| LHP030132 | MGB | Healthy | N/A | N/A | LPWGS | N/A | 0.00 | N/A | N/A |
| LHP030642 | MGB | Healthy | N/A | N/A | LPWGS | N/A | 0.00 | N/A | N/A |
| LHP031645 | MGB | Healthy | N/A | N/A | LPWGS | N/A | 0.00 | N/A | N/A |
| LHP034881 | MGB | Healthy | N/A | N/A | LPWGS | N/A | 0.00 | N/A | N/A |
| LHP035094 | MGB | Healthy | N/A | N/A | LPWGS | N/A | 0.00 | N/A | N/A |
| LHP038748 | MGB | Healthy | N/A | N/A | LPWGS | N/A | 0.00 | N/A | N/A |
| LHP041556 | MGB | Healthy | N/A | N/A | LPWGS | N/A | 0.00 | N/A | N/A |
| LHP056703 | MGB | Healthy | N/A | N/A | LPWGS | N/A | 0.00 | N/A | N/A |
| LHP098228 | MGB | Healthy | N/A | N/A | LPWGS | N/A | 0.00 | N/A | N/A |
